# Comprehensive characterization of transcript diversity at the human *NODAL* locus

**DOI:** 10.1101/254409

**Authors:** Scott D Findlay, Lynne-Marie Postovit

## Abstract

*NODAL*, a morphogen belonging to the transforming growth factor beta (TGβ) superfamily, is essential during embryogenesis where it induces axis formation and left-right asymmetry. *NODAL* is also required for the maintenance of human embryonic stem cell pluripotency, and emerges in many cancer types concomitant with metastasis and therapy resistance. Several enhancer elements have been shown to regulate mouse *Nodal* expression and studies have delineated mechanisms by which mRNA splicing and translation of NODAL homologues are regulated in model organisms. However, little is known regarding the co-transcriptional and post-transcriptional processing of human NODAL. Herein, we describe hitherto unreported RNAs which are transcribed from the *NODAL* locus, including an antisense transcript, a circular transcript, and multiple splice variants. These transcripts demonstrate the complexity of *NODAL* expression and highlight the need to consider each NODAL variant when attempting to quantify or target this morphogen.

## Introduction

The transforming growth factor-beta (TGF-β) superfamily member nodal growth differentiation factor (human gene symbol: *NODAL*, NCBI gene ID: 4838) plays essential roles in early embryonic development and is reactivated in cancers. *Nodal* is aptly named after its discovery in the mouse node in gastrula-stage embryos ^1^. *Nodal* has been well studied in numerous vertebrate embryos and *in vitro* models of early development ^2-5^. These studies have shown that *Nodal* promotes epiblast expansion and maintenance in blastocyst stage embryos ^6^, directs the specification of the distal visceral endoderm (DVE) after implantation ^7^, establishes the proximal-distal ^8^ and anterior-posterior ^8,9^ axes, influences mesendoderm differentiation during gastrulation (reviewed in ^5^), and establishes left-right asymmetry ^10^ ^11-13^.

Owing to practical and ethical limitations concerning research on human embryos, human-specific study of *NODAL* biology has generally been limited to cultured human embryonic stem (hES) cells where *NODAL* helps maintain pluripotency ^14^, and blocks differentiation toward neuroectoderm lineages ^15^. In addition, active SMAD2/3 downstream of *NODAL* (and other TGF-βs) also mediates cell fate decisions during mesendoderm differentiation ^16-19^.

*NODAL* expression in cancer was first identified by Postovit and Topczewska and colleagues in the aggressive C8161 human melanoma cell line ^20^. These cells were able to induce ectopic outgrowths or a complete secondary axis after injection into zebrafish embryos at the blastocyst stage and *NODAL* was identified as the primary factor responsible for this induction. Since this pioneering discovery, *NODAL* has been shown to affect numerous tumour phenotypes in experimental models of several human cancers including cancers of the breast ^21-25^, prostate ^26,27^, ovary ^28,29^, and pancreas ^30^, as well as glioma ^31,32^, glioblastoma ^33^, endometrial cancer ^34^, hepatocellular carcinoma ^35^, and choriocarcinoma ^24,36,37^. In these models, *NODAL* is generally pro-tumourigenic and impacts numerous processes including proliferation and apoptosis, migration and invasion, epithelial-to-mesenchymal transition (EMT), angiogenesis, and metastasis (reviewed in ^4^).

There is also strong correlative evidence of a link between high *NODAL* expression and poor clinical outcome in numerous cancers. For example, in a study of over 400 breast cancer patients, NODAL protein levels correlated positively with tumour stage and grade, independently of estrogen receptor/progesterone receptor (ER/PR) or HER2 status ^38^. Moreover, a recent meta-analysis of *NODAL* expression in human cancers originating from 11 different tissues and including more than 800 patients revealed significantly higher *NODAL* expression in cancerous tissue relative to healthy control tissue ^39^.

While transcriptional regulation of *Nodal* has been extensively studied in the mouse, and many enhancers have been identified ^10,40-48^, little is known regarding the co-transcriptional and post-transcriptional processing of NODAL transcripts. We recently reported how such regulation is relevant in our discovery of a genetically regulated alternatively spliced NODAL transcript in human pluripotent stem cells ^49^. This discovery, along with a previous report of inconsistent detection of NODAL transcripts when probing different exons ^50^, prompted us to perform a comprehensive quantitative analysis of human NODAL transcripts across human cell lines commonly employed to model *NODAL* biology. Herein, we describe multiple novel RNAs transcribed from the *NODAL* locus, including an antisense transcript, a circular transcript, and multiple splice variants. These RNAs can confound the interpretation of expression studies, particularly given variable levels of *NODAL* expression that can occur in response to different culture conditions.

## Results

Several studies have suggested that *NODAL* is essential for the maintenance of pluripotency in hES cells and a large-scale study demonstrated that *NODAL* and *NANOG* expression were highly correlated across numerous hES cell lines ^51^. However, few have thoroughly examined endogenous *NODAL* expression in response to various culture conditions. In our first attempts to use droplet digital PCR (ddPCR) assays for absolute quantification of NODAL in human embryonic stem cells, we made the surprising discovery that *NODAL* expression levels can vary dramatically in H9 hES cells. For example, one pair of samples differed in total NODAL transcript by over 3,000-fold, with only 26 copies of total NODAL transcript detected for the low-expressing sample in cDNA from 100 ng of total RNA (Fig. 1a,b).

**Figure 1:**
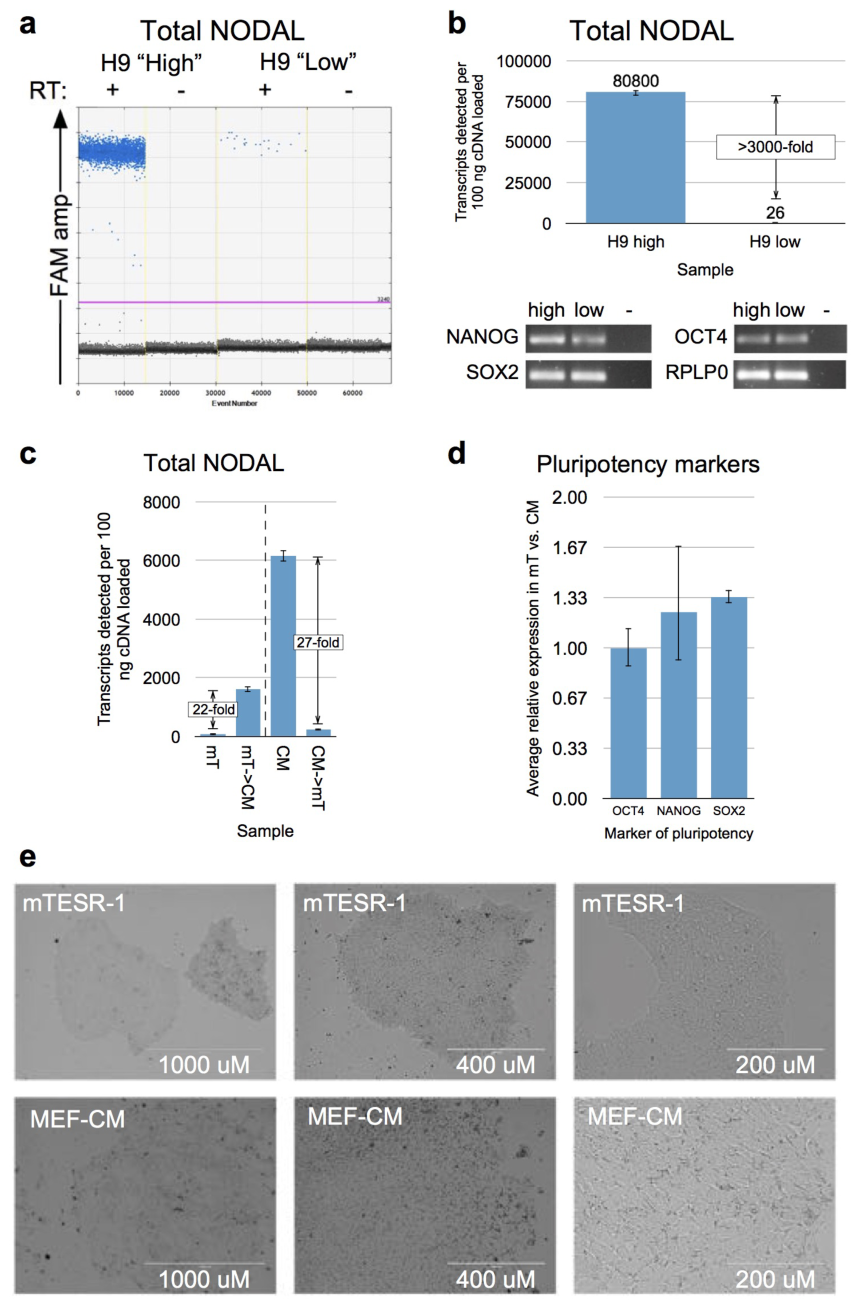
NODAL mRNA levels can vary dramatically between sub cultures of hES cells. a) ddPCR droplet plots for examples of “high” and “low” total NODAL expression in H9 hES cells using a primer probe assay spanning the exon 1 - exon 2 junction. “RT” = reverse transcriptase. Blue droplets are positive for NODAL target, and black droplets are negative. The pink line indicates amplitude threshold for a positive call. b) Top: Quantification of total NODAL levels from A. Bottom: Both “high” and “low” NODAL samples were positive for markers of pluripotency using end-point RT PCR. “-” = no template control. c) “mT” = H9 hES cells previously adapted to culture in defined mTESR-1. “CM” = H9 hES cells cultured in MEF-conditioned media for several passages. Arrows indicate transitions to different media. Error bars in b and c indicate 95% confidence interval for Poisson-calculated copies of transcript detected. d) Expression of pluripotency markers was not lower in mTESR-1 relative to MEF‐ CM culture conditions. Error bars indicate standard deviations. e) Representative images of hES cells cultured in mTESR-1 (top) and MEF-CM (bottom) at increasing magnifications (left to right). Scale bars are shown.

To experimentally investigate possible factors that may influence NODAL transcript levels, we focused on cell culture media, as either defined media such as mTESR-1, or media conditioned by mouse embryonic fibroblasts (MEFs), are regularly employed in the maintenance of hES cells. H9 hES cells adapted for culture in defined conditions (mTESR-1) expressed low levels of total NODAL transcript, but displayed markedly increased NODAL transcript levels after being switched to MEF-conditioned media (MEF-CM) for only several days (Fig. 1c). The reciprocal effect was also observed: when cells continually passaged in MEF-CM were returned to defined conditions, NODAL levels decreased by approximately the same factor as it had increased previously (Fig. 1c). Notably, cells under both conditions expressed similar levels of markers of pluripotency (Fig. 1d) and had morphologies typical of pluripotent stem cells (Fig. 1e). Hence, NODAL levels seem to fluctuate even when pluripotency is maintained.

We next sought to examine the exact composition of NODAL transcripts in human embryonic stem cells. We recently reported that a single nucleotide polymorphism (SNP) modulates a novel alternative splicing event for human NODAL in hES cells ^49^. Specifically, the minor allele of SNP rs2231947 in NODAL’s second intron contributes to splicing of a 116-base pair cassette alternative exon (Fig. 2). This splicing event was associated with both the sex of hES cell lines, as well as XIST RNA levels in female hES cell lines, but transcripts containing the alternatively spliced exon have yet to be fully characterized. Like total NODAL levels, the proportion of NODAL transcripts containing the alternative exon was also variable between sub cultures, and was higher in cells cultured in defined media relative to MEF-CM (Supplementary Fig. S1). For the remainder of this study, we aimed to always (when possible) distinguish the constitutively spliced (canonical) NODAL isoform from the newly discovered alternatively spliced NODAL variant isoform.

**Figure 2:**
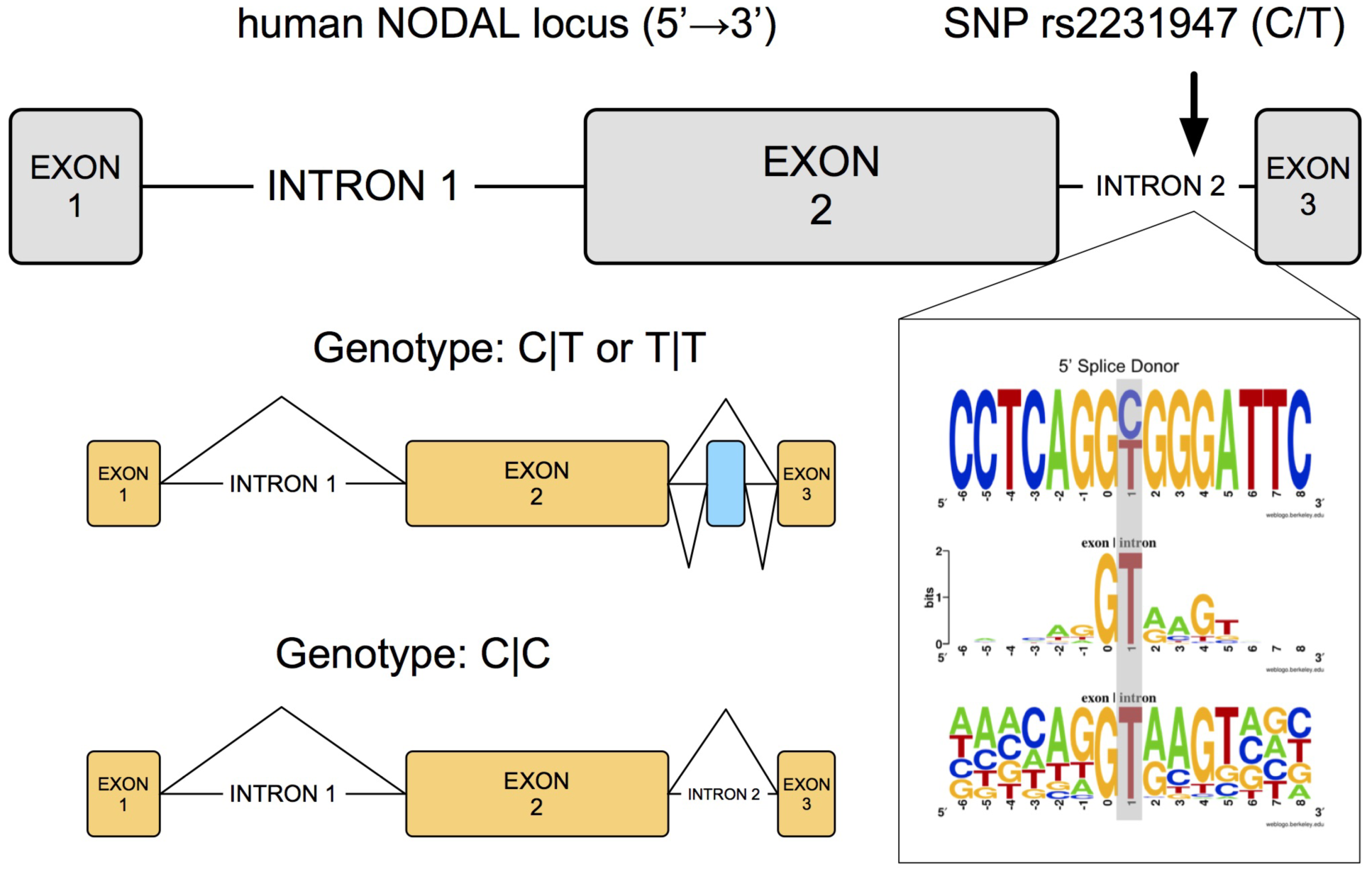
A genetically regulated splice variant is expressed by a subset of human pluripotent stem cells. Top: a schematic of the human *NODAL* gene is shown in grey. Diagram scale is approximate. Bottom left: the resulting splice variants are shown for different rs2231947 genotypes. Constitutively spliced exons are shown in yellow. The alternatively spliced exon is shown in blue. Bottom right: sequence “web logos” show human NODAL rs2231947 locus (top) relative to information content (middle) and nucleotide frequency (bottom) of human splice donor sites. Position “0” marks the exon-intron boundary and is the first (most 5’) base of an intron.

We first used rapid amplification of cDNA ends (RACE) to determine both the 5’ and 3’ termini. The 5’ end marks the transcriptional start site(s) and defines the length of the 5’ untranslated region (UTR) while the 3’ end defines the 3’ UTR and polyadenylation site(s) utilized. Standard 5’ RACE was conducted using a primer designed to reverse transcribe total NODAL, and a single product was obtained. Sequencing revealed a 5’ end 14 bases upstream of the annotated NODAL translational start codon, and 28 bases downstream of the annotated NODAL transcriptional start site in RefSeq (NM_018055.4). In contrast, several different products were detected for RNA reverse transcribed with a primer specific to the alternative NODAL exon (Supplementary Fig. S2). Two of these products revealed 5’ ends within the first exon of NODAL, but likely resulted from incompletely reverse-transcribed RNA. Interestingly, a third band revealed utilization of a novel upstream first exon spliced directly to exon 2. Subsequent analysis with RNA ligase mediated (RLM) 5’ RACE specific for capped mRNA ends confirmed a single major product for NODAL transcripts corresponding to transcriptional start at position −14 (Fig. 3a). A true 5’ end was also confirmed within the alternative upstream first exon using a separate primer set (Supplementary Fig. S2). However, it was not possible to *specifically* assess whether constitutively or alternative spliced NODAL isoforms contributed to any given 5’ end using the RLM method.

**Figure 3:**
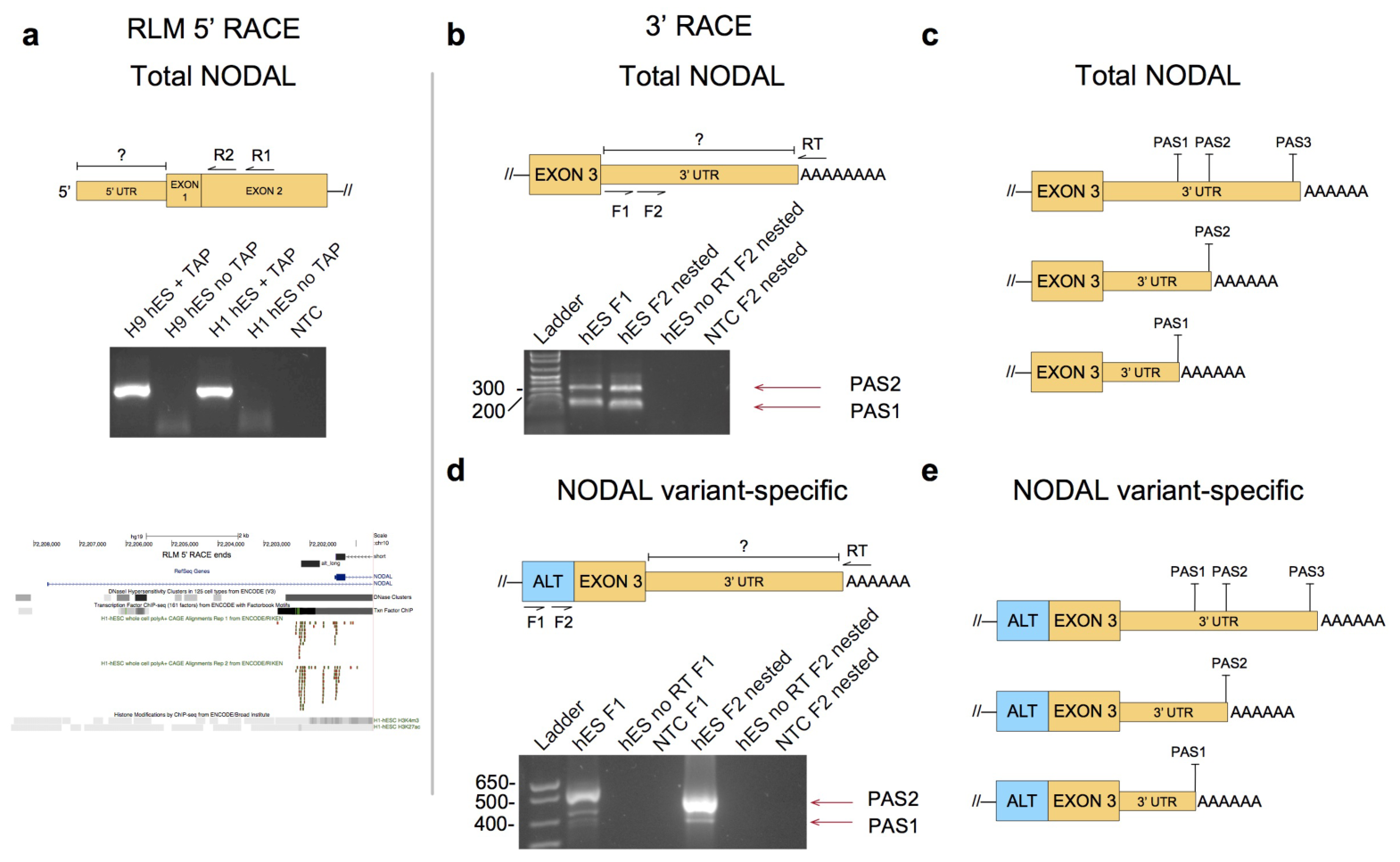
RLM 5’ RACE reveals a relatively uniform 5’ UTR, while 3’ RACE reveals alternative polyadenylation of NODAL transcripts. NODAL transcript schematics are shown at the top of each panel indicating the region of interest (“?”) and primers used for a, b, and d. “R1” = first reverse primer. “R2” = nested reverse primer. “F1” = first forward primer. “F2” = nested forward primer. “RT” = reverse transcription primer. “—//” indicates continuation of the transcript beyond what is shown. Adapter primer sites are not shown. a) Top: A single major 5’ end product was found for H9 and H1 hES cells. “TAP” = Tobacco Acid Pyrophosphatase. Bottom: Alignments of the major product 5’ end (“short”) and a minor product 5’ end (“alt_long,” see Supplementary Fig. S2) to the *NODAL* locus. Relevant tracks from the human genome browser are also shown, including Cap Analysis of Gene Expression (CAGE) read alignments that closely match both 5’ ends obtained in our RLM 5’ RACE analysis, and H3K4-trimethylation density in H1 hES cells. b) 3’ transcript end analysis reveals roughly equal utilization of two NODAL polyadenylation sites. c) These sites map to the two more proximal of three “canonical” polyadenylation sites (“PAS”) defined by A[A/T]TAAA motifs in the annotated NODAL 3’ UTR. d) and e) The same analysis reveals polyadenylation of NODAL variant transcripts at the same two sites, but with usage skewed heavily toward the more distal site. Constitutive exons are shown in yellow and the alternative cassette exon (“ALT”) is shown in blue. “AAAA…” represents the polyA tail at the 3’ end of transcripts. Exons upstream (5’) of the alternative exon or exon 3 are not shown.

For processed mRNA transcripts, 3’ ends are marked by the start of a polyA tail approximately 15-30 nucleotides downstream of a polyadenylation signal (PAS) (reviewed in ^52^). Analysis of NODAL’s constitutive terminal exon 3 for common polyadenylation signals revealed two AUUAAA motifs and a single AAUAAA motif (Fig. 3c). These two motifs are the most commonly utilized for polyadenylation of human transcripts ^53^, although other less-frequently utilized putative PASs were also found in the annotated 3’ UTR. 3’ RACE for total NODAL transcript revealed roughly equal utilization of either a more proximal AUUAAA site, or a more distal AAUAAA site (Fig. 3b). NODAL variant transcripts also utilized the same polyadenylation sites, but in a manner highly skewed toward the distal site (Fig. 3d).

We were next interested in comparing the performance of assays targeting different regions of the NODAL transcript. Since ddPCR is absolutely quantitative, it permitted direct comparison of the abundance of different targets and was selected as our PCR platform. For the same “high NODAL” sample from Fig. 1, the number of transcripts detected did not differ substantially when targeting different regions of the transcript: There was less than a 1.5-fold difference seen when targeting the exon 2 - exon 3 junction, exon 1 – exon 2 junction, or exon 2 only (Fig. 4a). However, for the “low NODAL” H9 sample, signal was much higher when probing exon 2 only, relative to either of the exon junctions (Fig. 4a). “No reverse transcription” controls demonstrated this signal was specific to RNA and did not result from genomic DNA or other DNA contamination (Supplementary Fig. S3). The lack of increased signal within exon 2 for the “high NODAL” sample suggests that exon 2 is not more efficiently reverse transcribed, but instead that additional transcript(s) sharing sequence with exon 2 may exist. Unless co-regulated, these would make a higher relative contribution to total signal when NODAL levels are low. A survey of human RNAs from Genbank ^54^ revealed AK001176 as a candidate transcript that completely encompasses exon 2 of *NODAL* (Fig. 4b). We developed a ddPCR assay specific to AK001176 unable to detect spliced full-length NODAL and found that polyadenylated AK001176 was expressed in hES cells (Fig. 4c). AK001176 was transcribed in the antisense direction relative to full-length NODAL and can therefore be classified as a natural antisense transcript (NAT) to NODAL (Supplementary Fig. S4). Interestingly, AK001176 was also alternatively polyadenylated and contains an open reading frame that may code for a protein (Supplementary Fig. S4).

**Figure 4:**
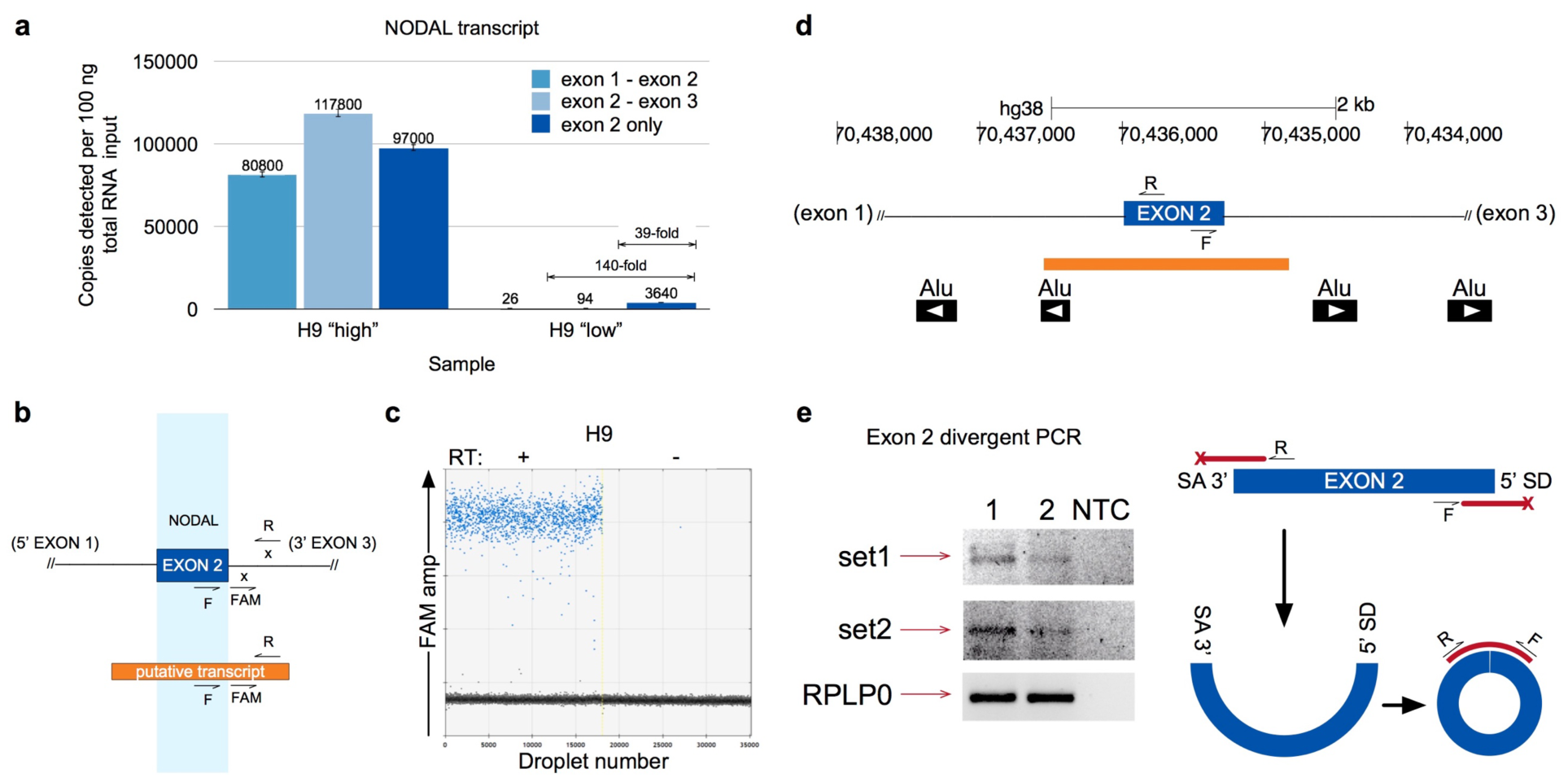
A natural antisense transcript (NAT) and circular RNA share sequence with and confound detection of NODAL’s second exon. a) NODAL detection across different regions of the full-length transcript. b) Design of ddPCR assay to specifically detect the NAT. c) Quantification of NAT. Blue dots represent positive droplets. Black dots represent negative droplets. “RT” = reverse transcriptase. d) Locations of Alu SINE elements are shown relative to the *NODAL* gene. Arrows indicate orientation/strand of Alu elements. e) Left: End-point PCR detection of circular exon 2 amplicons (and products resulting from template switching) using two different primer sets in two different H9 hES samples (“1” and “2”). “NTC” = no template control. Images were inverted for better visualization of bands. Right: Schematic of NODAL exon 2 circular RNA and PCR strategy used. Red bars indicate PCR amplicons. “x” indicates non-productive amplification of linearly-spliced exon 2. “SA” = splice acceptor. “SD” = splice donor. For all schematics, NODAL exon 2 is shown in blue and the NAT in orange. “F” = forward primer. “R” = reverse primer. “x” indicates no primer binding site in full-length NODAL transcripts. “FAM” = fluorescent probe.

We were also interested in testing for the potential presence of other transcripts containing exon 2 sequence and discovered that NODAL exon 2 was an excellent candidate to form a circular RNA. Circular RNA occurs when the 5’ splice donor site of an intron forms a “back splice” with an upstream 3’ splice acceptor site of the same or other exons in the transcript ^55^. Relative to splice sites in general, it has been shown that circular RNA splice sites are more likely to be flanked by upstream and downstream intronic Alu repeat elements and that these genomic elements are more likely to be in opposite orientations. Single circularized exons were also found to be formed from some of the longest of all human exons, with an average length of 690 nucleotides ^55^. In addition to constitutive exon 2 of *NODAL* being an extremely long exon (698 nucleotides), there are two pairs of Alu repeats in opposite orientations in the intronic sequences flanking *NODAL* exon 2 (Fig. 4d). We used divergent PCR to determine if a circular RNA was present and detected a single circular RNA product formed by back splicing of exon 2 of NODAL (Fig. 4e).

Having now detected novel antisense and circular transcripts in addition to our previous discovery of an alternatively spliced transcript, we were motivated to profile all of these *NODAL* locus RNAs across several human cell lines that have been used to model *NODAL* biology. To this end, we assembled a panel that included breast cancer cell lines (both Estrogen Receptor positive and negative), where *NODAL* has been extensively modelled ^22-24,56^, the C8161 melanoma cell line where NODAL was first characterized in cancer ^20^, HEK 293 cells of embryonic origin, and H9 hES cells (cultured in MEF-CM). All regions of full-length NODAL and all of the newly discovered transcripts were profiled in parallel using ddPCR. For all cancer cell lines assayed, extremely low (or undetectable) levels of NODAL transcript (≤ 2 copies per 100 ng total RNA input) were detected using assays spanning either exon-exon boundary. In contrast, signal detected within exon 2 ranged from 316 to 1,786 copies of transcript per 100 ng total RNA input (Fig. 5). However, this assay is not specific for full-length NODAL and also detects the AK001176 NAT. Indeed, very similar levels of transcript were detected for AK001176 and the NODAL exon 2 assay across all cancer cell lines. Specifically, the maximum ratio of NODAL exon 2 signal to AK001176 signal among cancer cell lines was 1.6-to-1, with an average of 1.4-to-1. These two signals were also highly correlated, with variability in AK001176 signal explaining 96% of the variability in exon 2 signal among cancer cell lines (Fig. 5b). Similar levels of AK001176 transcript were detected between H9 hES and cancer cells, while the circular NODAL RNA was detected only in H9 hES cells. Collectively, these results suggest that bulk cancer cell lines express very little processed (spliced) NODAL transcript, and that use of assays confined to exon 2 is inappropriate for the detection of full-length NODAL, owing to consistent expression of a *NODAL* NAT containing sequence identical to the entirety of *NODAL* exon 2.

**Figure 5:**
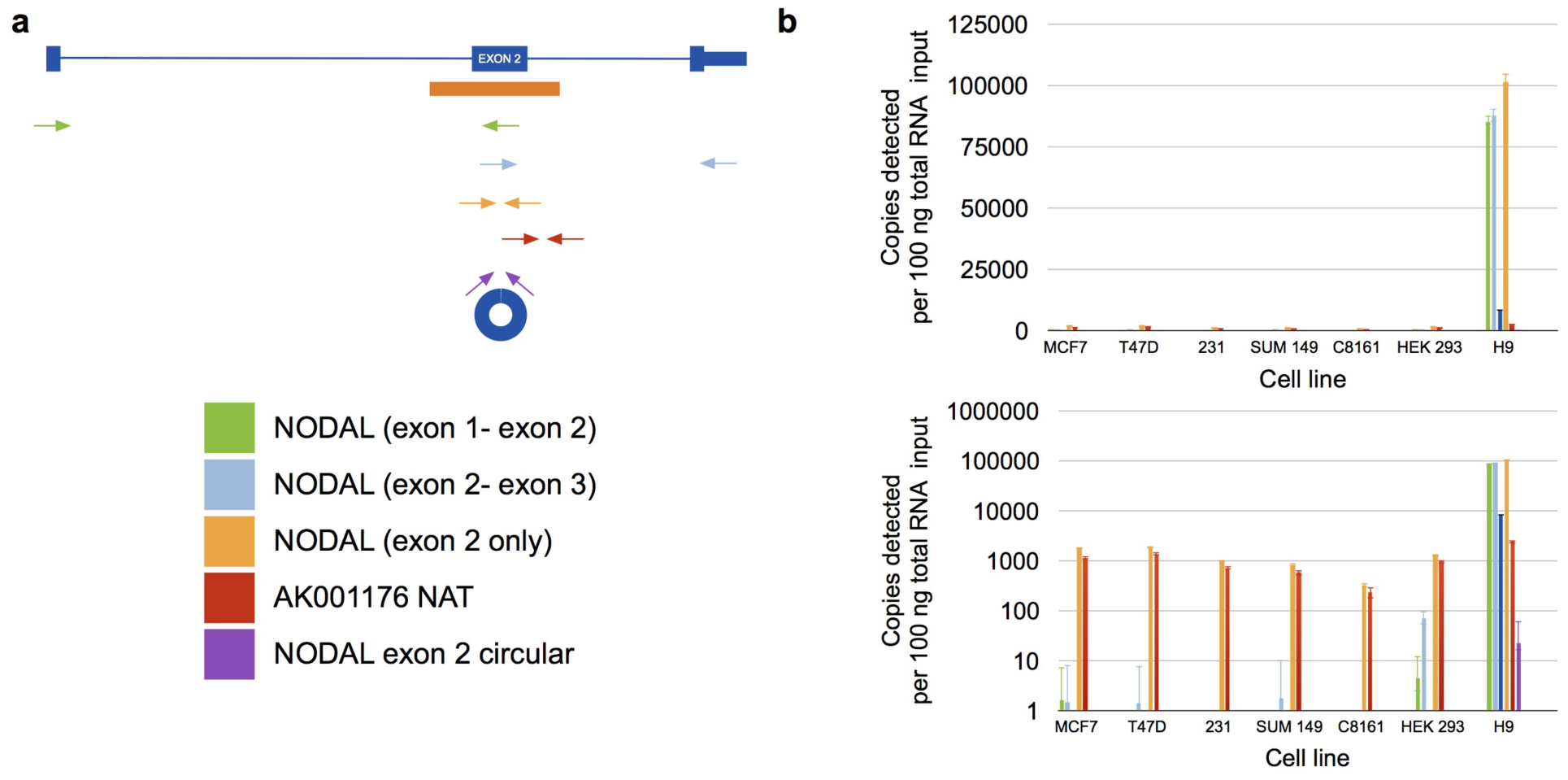
Profiling of *NODAL* locus transcripts using ddPCR. a) Approximate target location for ddPCR assays used to quantify transcripts expressed from the *NODAL* locus. Full-length NODAL is shown in blue, and the NAT in orange. Arrows indicate approximate location of forward and reverse primers only. For simplicity, locations of fluorescent probes are not shown. b) Quantification of all *NODAL* locus transcripts in parallel. The bottom chart illustrates the same data as the top, but plotted on a log10 scale. “231” = MDA-MB-231. Error bars indicate 95% confidence intervals of Poisson-calculated target copies.

## Discussion

In this work, we have characterized and quantified both known and novel transcript isoforms expressed from the human *NODAL* locus in cell lines where this gene is commonly studied. Specifically, in addition to a previously identified genetically-regulated splice variant ^49^, we detected an alternative transcriptional start site and first exon upstream of constitutive exon 1, alternative polyadenylation site usage, a circular RNA formed by back-splicing of exon 2, and a natural antisense transcript encompassing the exon 2 locus. Collectively, these results point to complex regulation of *NODAL* gene expression at the RNA level. The discoveries reported here should be used to refine assay design for more specific and accurate NODAL detection, and to improve strategies designed to experimentally (and potentially therapeutically) target *NODAL*.

We also characterized multiple aspects of the full-length NODAL transcripts, discovering fairly uniform short 5’ UTR usage (although a rare alternative 5’ UTR was also detected), and alternative polyadenylation at two PAS sites in the 3’ UTR. These aspects of NODAL processing present interesting opportunities to advance understanding of NODAL regulation, as both alternative transcriptional start site (TSS) utilization as marked by the 5’ UTR, and alternative polyadenylation (APA) marked by the 3’ UTR can influence mRNA dynamics including translation ^57-59^. Furthermore, global changes in 3’ UTR length resulting from APA occur in both early embryonic development ^60-62^ and in oncogenesis ^63-65^—both highly relevant cell contexts for NODAL.

The apparent enrichment of an alternative first exon and skewed polyadenylation site usage for the *NODAL* variant relative to constitutively spliced *NODAL* suggests there may be coordinated regulation of the *NODAL* variant transcript that extends beyond the splice donor site formed by the rs2231947 T allele in cis (Fig. 2 and Fig. 3). However, this apparent enrichment may also result from technical issues such as differential reverse transcription efficiencies of distinct transcripts. The apparent coordination with polyadenylation is interesting given that a link between alternative polyadenylation and alternative splicing has been described, but only for 3’ terminal exon selection and intronic polyadenylation sites ^66,67^.

The ability to quantitatively profile expression levels of all *NODAL* locus RNAs was enhanced by ddPCR. As an absolutely quantitative method, resultant transcript levels are not influenced by PCR efficiencies or dependent on the use of standard curves. Furthermore, duplexed ddPCR assays were used to easily distinguish constitutively spliced NODAL transcripts from those containing the alternative cassette exon (Supplementary Fig. S1).

Our finding that NODAL transcripts were not consistently highly expressed in hES cells was surprising given that *NODAL* signalling is generally thought to be essential for hES cell pluripotency ^14,17^ and consistently high NODAL expression has been reported for this cell type ^51^. Our “low NODAL” cells maintained typical pluripotent stem cell morphology and expression of markers of pluripotency (Fig. 1e). It is possible that low NODAL levels are indicative of suboptimal cultures poised for (or already undergoing) early differentiation. In this vein, NODAL, LEFTY1, and LEFTY2 displayed some of the most rapid down-regulation upon spontaneous hES cell differentiation in a small panel of pluripotency markers ^68^. It is also possible that high NODAL mRNA expression is not strictly required for the maintenance of pluripotency, and that NODAL is either preferentially translated into protein, or that there are redundant or compensatory mechanisms that can otherwise sustain pluripotency in certain culture conditions when NODAL mRNA levels are low. Our results are consistent with the notion that a pluripotent gene expression signature is not static or universal, but rather partially stochastic, and that the combinations of active transcription factor networks and signalling pathways that can support the pluripotent state can drift with culture conditions and microenvironmental factors, between cell lines, and due to other unknown variables. Indeed, it has been suggested that there exists a spectrum or continuum of pluripotent states both *in vitro* and *in vivo* (reviewed by ^69^). In our study, the observed variability in hES cell NODAL transcript levels was certainly staggering. That NODAL transcript levels were dramatically and reversibly influenced by culture conditions may be an indication that more general differences exist between hES cells cultured in MEF-conditioned serum replacement-based media and in more defined media. In general, some differences in hES cells cultured under varying conditions have been observed ^70,71^. However, very little work has directly compared culture of hES cells in MEF-CM to culture in defined media such as mTESR ^72^. Perhaps surprisingly, there is a striking absence of work involving comprehensive and quantitative profiling of hES gene expression signatures between MEF-CM and defined media, to investigate to what extent such media may potentiate distinct pluripotent states *in vitro*.

In addition to the unexpectedly low levels of NODAL transcript sometimes observed in hES cells, we also made the surprising discovery of especially low *NODAL* expression at the transcript level across numerous human cancer cell lines. The extremely low levels and often absence of any NODAL transcript reported here for cell lines such as C8161 aggressive melanoma and MDAMB-231 triple-negative breast cancer are inconsistent with functional studies in these cell lines where NODAL knockdown, mediated through either RNA interference ^22,25^, or inhibition of endogenously expressed protein ^73^ resulted in profound phenotypic effects. Notably, NODAL mRNA expression was not assessed in these papers, so it is difficult to tell whether the detectable and functionally relevant levels of NODAL protein reported in these studies were expressed from cells with considerably higher NODAL mRNA levels. There are several possibilities for this apparent discrepancy. First, it is possible that NODAL mRNA is preferentially stabilized or translated, generating high levels of protein from a limited transcript pool. Second, it is possible that a technical issue such as NODAL-specific extremely inefficient reverse transcription limits the detection of transcript, although it is especially important to emphasize that high levels of NODAL transcript were routinely detected in hES cell samples analyzed in parallel. Third, it is possible that NODAL mRNA steady state levels are highly heterogeneous between subcultures of cancer cell lines or that this transcript is only present in a very rare subpopulation (for example cancer stem cells) such that the levels are diluted in bulk cultures. Finally, in zebrafish, mRNA of the maternal *Nodal* homolog *sqt* is prevalent as unspliced pre-mRNA ^74-76^. It is possible that cancer cells similarly retain NODAL mRNAs in this unprocessed state, which would not be detectable using primers spanning introns. Regardless, the quantitative profiling performed here certainly helps explain some of the previously reported challenges associated with detection and modelling of *NODAL* in human cell lines ^50^.

Consistent with our findings, at least one other group has reported undetectable *NODAL* transcript levels in MDA-MB-231 cells when using a real time PCR assay spanning an exon-exon boundary

^47^. One other study has directly compared NODAL expression levels between two cancer cell lines and hES cells with multiple assays, although this analysis was conducted using semi-quantitative end-point PCR ^50^. For the C8161 cell line, an assay crossing the exon 2 - exon 3 boundary resulted in a low intensity band. In contrast, an assay internal to exon 2 yielded a band of much higher intensity. This result is consistent with those presented here, which revealed that this higher signal may at least partially result from confounded detection of the AK001176 NAT sharing sequence with exon 2 of *NODAL*. The increased signal from assays internal to exon 2 is likely not the result of higher reverse transcription efficiency in this region of the transcript since signal was fairly uniform across all regions of the transcript in an H9 sample with high *NODAL* expression. A major conclusion of this work is that assays targeting only exon 2 of NODAL are not specific to full-length spliced NODAL transcripts and are thus inappropriate for NODAL transcript detection. Unfortunately, such assays have already been widely employed when assessing NODAL mRNA levels (e.g. ^22,26,30,50,56,77-80^). Going forward, it is highly recommended that specific assays for the antisense transcript, circular RNA, and NODAL splice variants be employed to untangle the contributions of each transcript to any overall change in expression measured by the assays internal to exon 2, as conducted in Fig. 5. As an example, it will be interesting for future studies to explore whether these three transcripts show similar responses to altered microenvironments.

In conclusion, we believe the *NODAL* locus is an interesting example of the capacity the cell has for complex RNA transcription and processing. There is reason to believe that *NODAL* is more the rule than the exception in this regard, as genome-wide analyses of processes such as alternative splicing and alternative polyadenylation have revealed multiple transcripts for the vast majority of protein coding genes ^81-84^. Still, the full-length nature and functional consequences of many alternatively processed transcripts are only beginning to be appreciated on a genome wide scale ^85^. Of course, the alternative processing of RNAs has major impacts on either the sequence or levels of resulting protein, which in turn affect cellular function and phenotypes. In order to advance our general understanding of gene expression and function, we must move beyond the simple conceptualization of a single RNA corresponding to each gene, and fully explore the staggeringly complex intracellular world at the RNA level.

## Methods

### Web logos

Sequence logos in Figure 2 were created using WebLogo version 2.8.2 (http://weblogo.berkeley.edu/ and ^86,87^).

### Microscopy

Pictures of H9 hES cells in different media were taken using EVOS FL Cell Imaging System (Thermo Fisher) with either 4X, 10X, or 20X objective lenses. Contrast and other image properties were adjusted so that cells and colony boundaries were more easily visible.

### Cell culture

H1 and H9 lines were purchased from WiCell (Madison, Wisconsin, USA). H9 cells were maintained on irradiated CF-1 Mouse Embryonic Fibroblasts (MEFs) (GlobalStem; Gaithersburg, Maryland, USA) with standard media composed of DMEM/F-12 with GlutaMAX^TM^ (Thermo Fisher Scientific; Waltham, Massachusetts, USA), 20% KnockOut^TM^ serum replacement (Thermo Fisher), 1X non-essential amino acids (Thermo Fisher), 0.1 mM 2-mercaptoethanol (Thermo Fisher), and 4 ng/ml of basic fibroblast growth factor (Thermo Fisher). H1 cells were maintained in feeder-free conditions that consisted of growth on a Geltrex matrix (Thermo Fisher) with either defined mTeSR1 media (Stem Cell Technologies; Vancouver, British Columbia, Canada), or CF-1 MEF-conditioned media as indicated. All hES cells were passaged manually and always harvested from feeder-free culture conditions. HEK 293 (ATCC; Manassas, Virginia, USA) cells were maintained in DMEM (Thermo Fisher) supplemented with 10% fetal bovine serum (Thermo Fisher). T47D, MCF7, and MDA-MB-231 breast cancer cells, and C8161 melanoma cells ^88^, were cultured in RPMI supplemented with 10% FBS (Thermo Fisher). SUM149 cells were cultured in media consisting of Ham’s F12 with 5% heat inactivated FBS, supplemented with HEPES, Hydrocortisone, and Insulin according to instructions provided by Asterand Bioscience (Detroit, USA). All cells were cultured at 37°C with 5% C0_2_ supplementation in a humidified environment.

### RNA extraction

Total RNA was isolated from cultured cells using the PerfectPure RNA Cultured Cell Kit (5-Prime; Hilden, Germany) or the RNeasy mini kit (Qiagen; Hilden, Germany), including on-column DNase treatment, and quantified with the Epoch plate reader (Biotek; Winooski, Vermont, USA).

### cDNA synthesis (for non-RACE analyses)

Total RNA was reverse transcribed with the high capacity cDNA reverse transcription kit (Applied Biosystems; Foster City, California, USA) following manufacturer’s instructions. Random hexamers were generally used to prime synthesis by reverse transcriptase. Reactions where oligo dT was used in place of random hexameters are indicated. “No RT” control reactions included RNA template and all components except reverse transcriptase enzyme. Figures with “transcripts detected per x ng cDNA loaded” axes refer to the amount of cDNA used based on the quantification of RNA. A separate quantification of cDNA was not performed.

### Droplet digital PCR (ddPCR)

ddPCR for total NODAL was conducted using Taqman primer probe assays Hs00415443_m1 (exon 1 – exon 2), Hs00250630_s1 (exon 2 only), or Hs01086749_m1 (exon 2 – exon 3) (Applied Biosystems). Where not indicated, Hs00415443_m1 (exon 1 – exon 2) was used for detection of total NODAL transcript.

The following primers and probes were used for duplexed detection of NODAL splice variants, with fluorophores, internal quenchers, and terminal quenchers flanked by forward slashes.

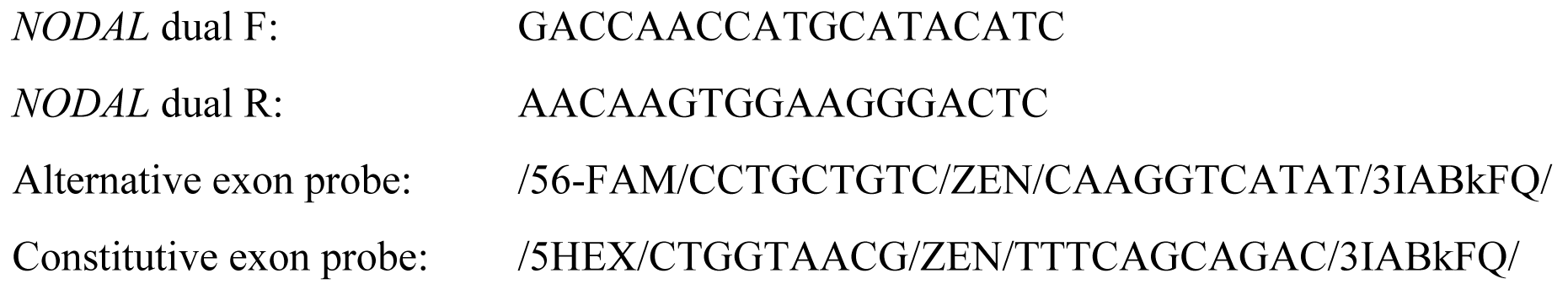

Droplets that were both FAM^+^ and HEX^+^, corresponding to the *NODAL* variant, were quantified using Quantasoft software. Since constitutive *NODAL* was FAM^-^ and HEX^+^, and could therefore be co-amplified in droplets containing *NODAL* variant transcript, constitutive *NODAL* was calculated manually using the equation: copies/ 20 μL sample = ‐ln(1-p) x 20,000 / 0.85. where ‘p’ is the proportion of positive droplets defined as FAM^-^HEX^+^ droplets / (FAM^-^HEX^+^ droplets + empty droplets), and 0.85 nL is the average volume of a droplet as used by QuantaSoft (Bio-rad)^89^.

For detection of the *NODAL* NAT transcript, the following primers and probe were used:

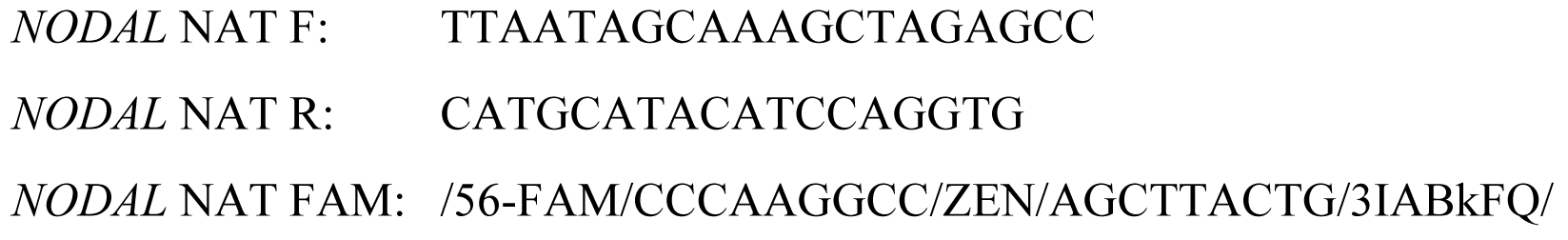

For detection of the *NODAL* circular exon 2 transcript, the following primers and probe were used:

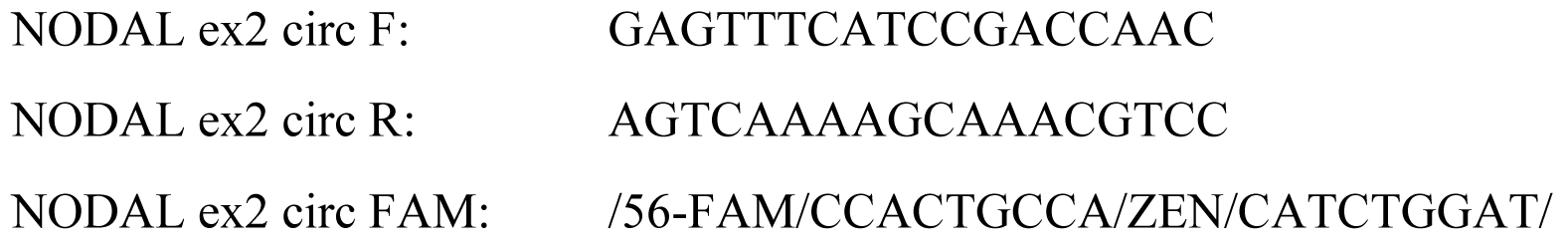

For all ddPCR assays, primers were used at a final concentration of 900 nM and probes were used at a final concentration of 250 nM.

For the dual NODAL splice variant assay, a “two-step” PCR was used with the following conditions:

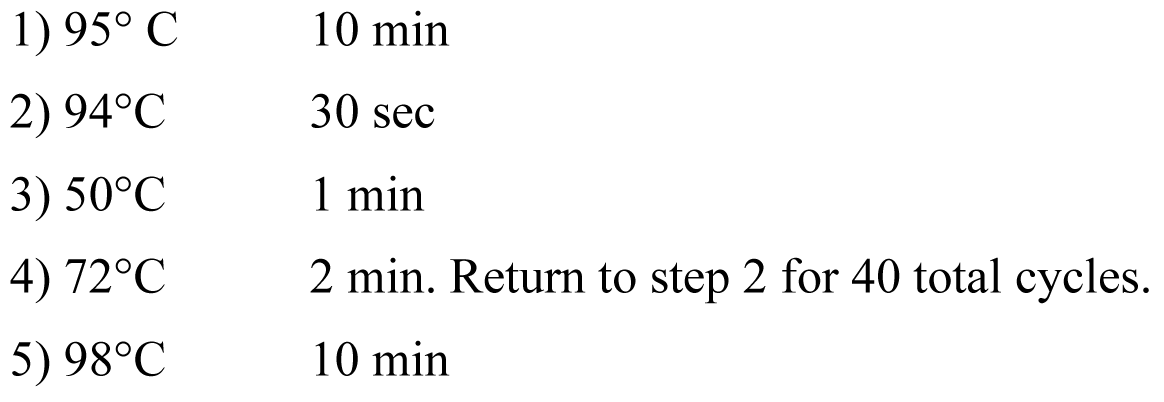

For all other ddPCR assays, the following cycling conditions were used:

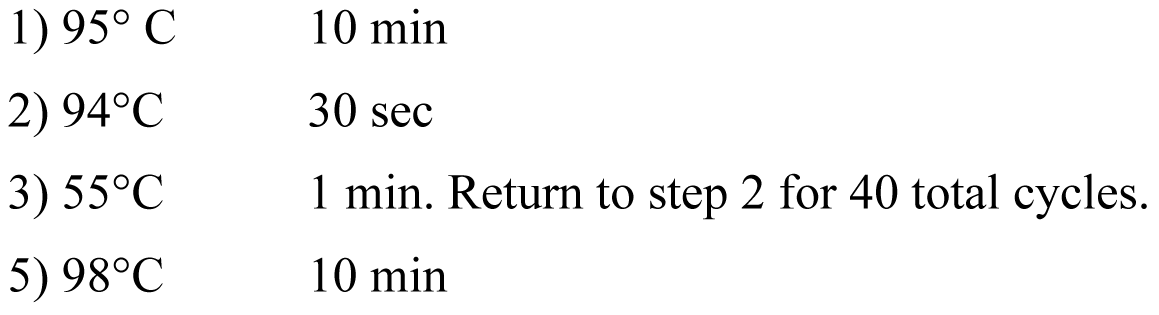

### Real time PCR

Real time PCR was performed using Taqman gene expression master mix (Applied Biosystems) and Taqman gene expression assays for POU5F1 (also known as OCT4) (Hs04260367_gH), NANOG (Hs04260366_g1), SOX2 (Hs01053049_s1), RPLP0 (4333761), and TBP (Hs99999910_m1). Expression was normalized to both RPLP0 and TBP using the ∆∆Ct method.

### RNA ligase mediated (RLM) 5’ RACE

RLM 5’ RACE was performed with the FirstChoice RLM-RACE kit (Ambion) according to manufacturer’s instructions. 10 μg of total RNA from H1 or H9 hES cells was used. Reverse transcription was performed at 50°C for one hour using random decamers provided. No-TAP, no-RT, and no-template control reactions were included to ensure specificity of products obtained. Subsequent PCR reactions were performed with AmpliTaq Gold 360 Master Mix with 1 μL of cDNA per reaction. An annealing temperature of 55°C was used for all reactions.

The following primers were used:

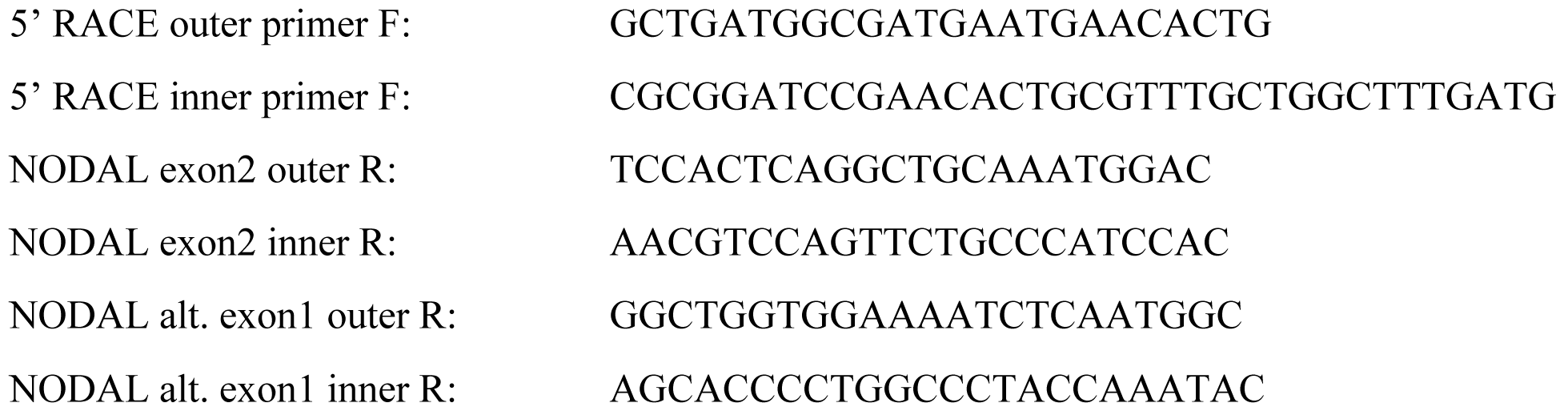

### Standard 5’ RACE

Standard 5’ RACE analysis was conducted using the 5’ RACE System for Rapid Amplification of cDNA Ends (Thermo Fisher) following manufacturer’s instructions. Three (3) μg of total RNA was used for each sample. Reverse transcription was performed for 50 minutes. All primers for first and second round PCR were used at a final concentration of 400 nM. An annealing temperature of 56°C was used for all reactions.

Primers used for reverse transcription:

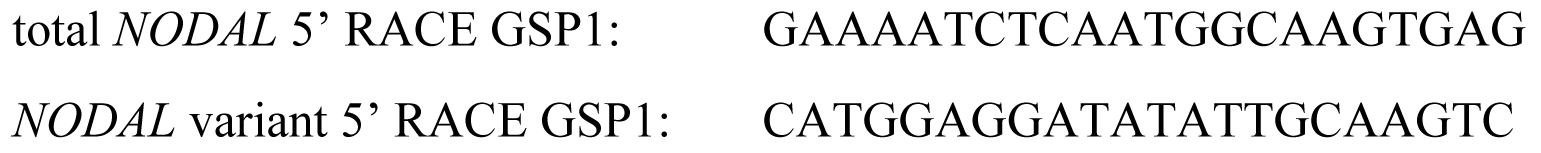

Primers used for first round PCR:

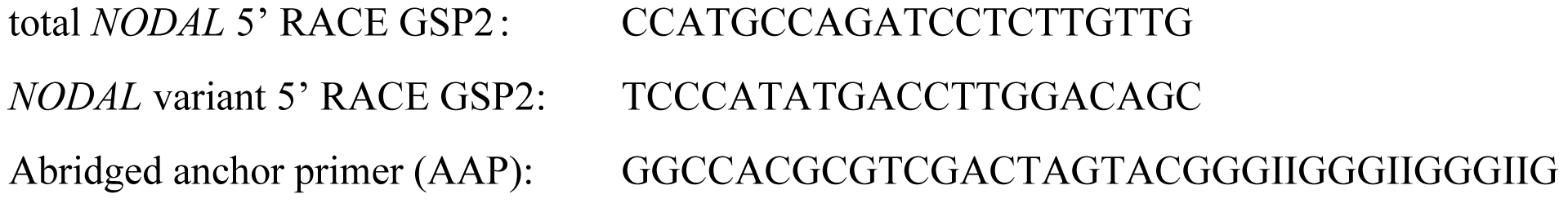

The same primer targeting constitutive exon 2 of *NODAL* was used for second round nested PCR analysis of both total *NODAL* and *NODAL* variant transcripts:

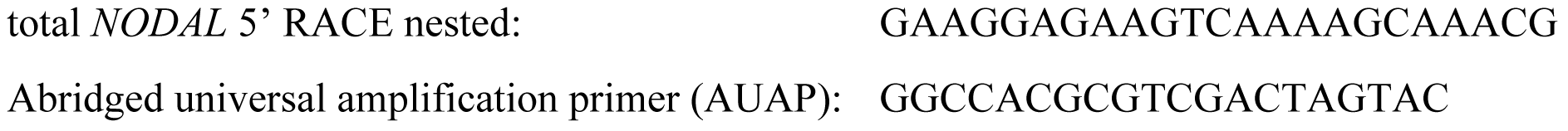

### 3’ RACE

For 3’ RACE, 2 μg total RNA was used for reverse transcription. Random primers were substituted for an oligo dT-adapter mix of “lock-dock” ^90^ primers with either A, G, or C as the most 3’ base:

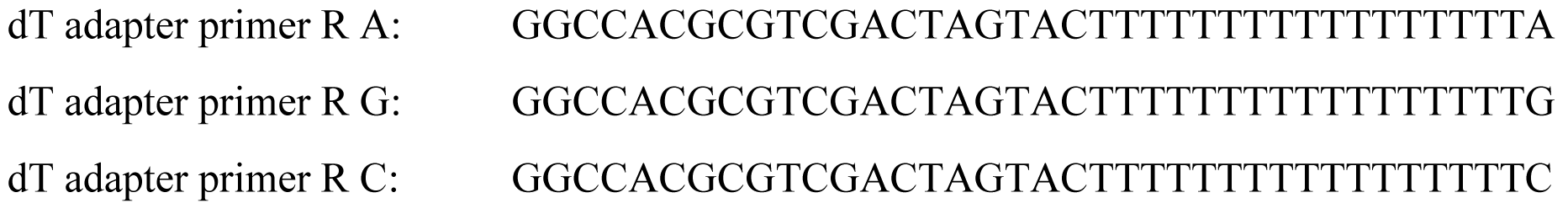

Each primer was used at a final concentration of 167 nM for a total primer concentration of 500 nM. 2 μL (equivalent to 200 ng RNA) of each cDNA reaction was used for subsequent PCR performed with AmpliTaq Gold 360 Master Mix. Primers were used at a final concentration of 200 nM. An annealing temperature of 54°C was used for all reactions.

Forward primers (variable for each target):

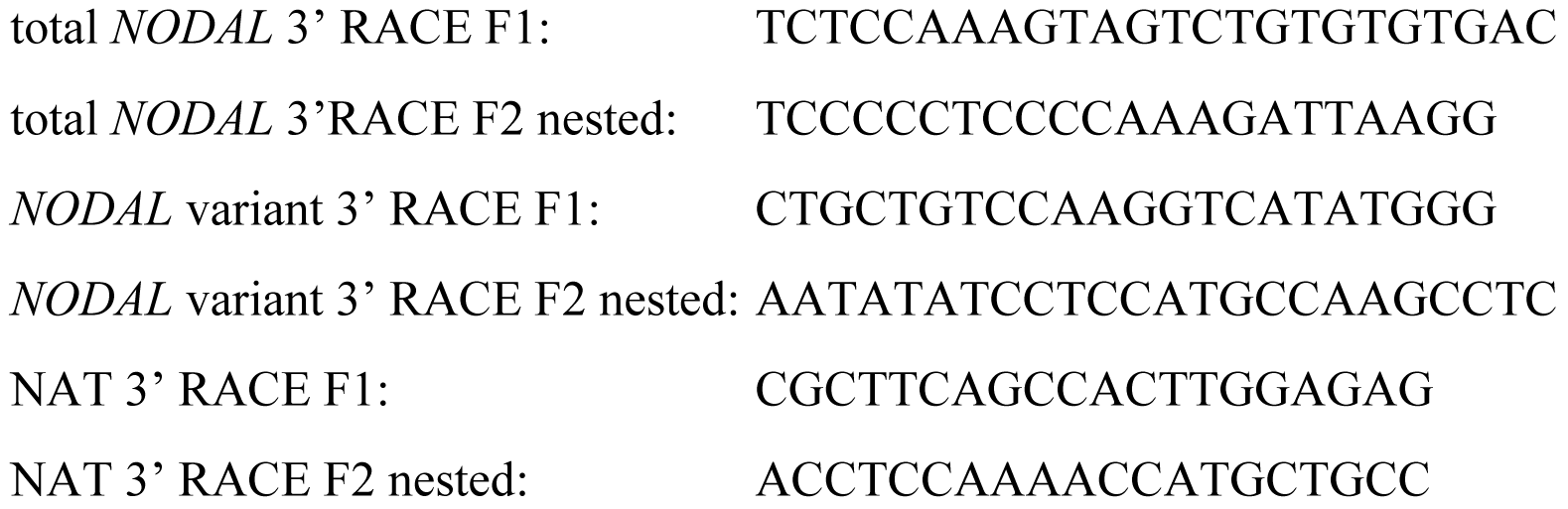

Reverse primer (identical for each target):

Abridged universal amplification primer (AUAP) R: GGCCACGCGTCGACTAGTAC

### Other (non-RACE) end-point PCR

AmpliTaq Gold 360 Master Mix (Applied Biosystems) was used for all end-point PCR analyses. Primers were used at a final concentration of 250 nM.

For the *NODAL* exon 2 divergent PCR to detect circular RNA, the following primers were used:

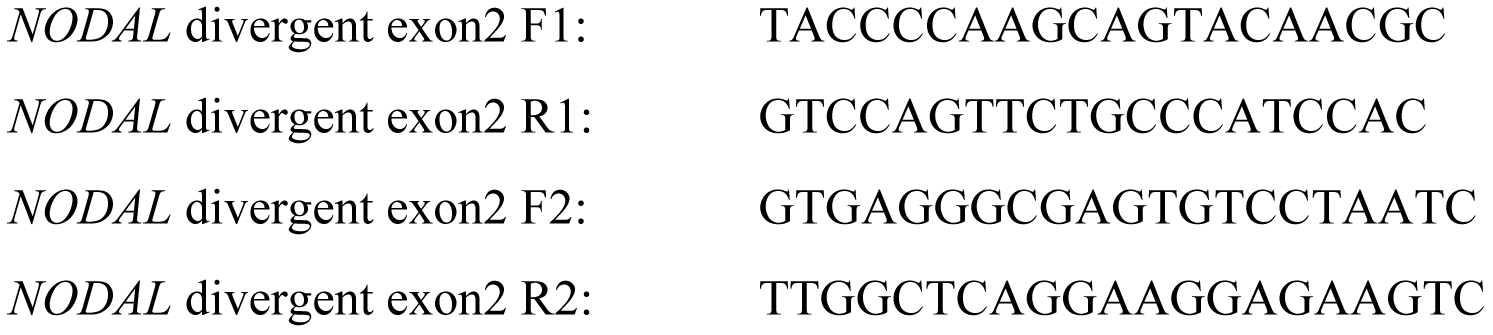

An annealing temperature of 55°C and an extension time of 1 minute were used.

To detect the *NODAL* NAT, the following primers were used:

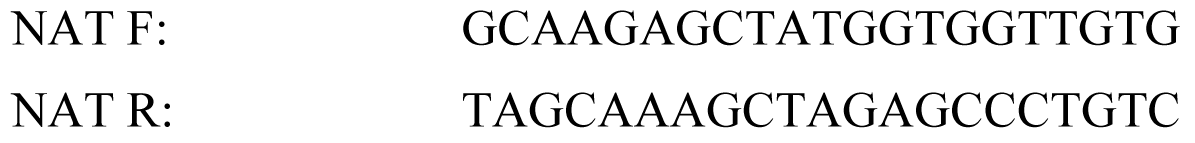

An annealing temperature of 54°C and an extension time of 2 minutes were used.

### Cloning and sequence analysis

All RACE and other end-point PCR products amplified with custom primers were cloned into the pCR 4-TOPO plasmid with TOPO TA cloning for sequencing kit (Thermo Fisher). Cloning reactions were transformed into One Shot TOP10 Chemically Competent *E. coli* (Thermo Fisher). Individual clones were selected with Kanamycin and propagated for mini prep of plasmid DNA using the High-Speed Plasmid Mini Kit (Geneaid/FroggaBio; Toronto, Ontario, Canada). Multiple clones were sequenced for each product to confirm amplicon identities. Sanger sequencing using the plasmid-specific M13R or M13F primers was conducted by the Molecular Biology Service Unit at the University of Alberta (Edmonton, Canada), or the London Regional Genomics Centre at Western University (London, Canada).

## Acknowledgements

This work was supported by an Alberta Innovates Health Solutions Translational Health Chair in cancer and by the Sawin-Baldwin Chair and the Dr. Anthony Noujaim chair from WCHRI and the ACF awarded to LMP.

## Author Contribution Statement

Conceived and designed the experiments: SDF and LMP. Performed the experiments: SDF. Analyzed the data: SDF. Wrote the paper: SDF and LMP. Reviewed and approved the final manuscript: SDF and LMP.

